# Interspecies interactions determine growth dynamics of biopolymer degrading populations in microbial communities

**DOI:** 10.1101/2023.03.22.533748

**Authors:** Glen D’Souza, Julia Schwartzman, Johannes Keegstra, Jeremy E Schreier, Michael Daniels, Otto Cordero, Roman Stocker, Martin Ackermann

**Author notes:** Corresponding author: Glen G D’Souza, Address: Eawag, Ueberlandstrasse 133, 8600 Duebendorf, Switzerland.

## Abstract

Microbial communities perform essential ecosystem functions such as the remineralization of organic carbon that exists as biopolymers. The first step in mineralization is performed by biopolymer degraders, which harbor enzymes that can break down polymers into constituent oligo- or monomeric forms. The released nutrients not only allow degraders to grow, but also promote growth of cells that either consume the breakdown products, i.e., exploiters, or consume metabolites released by the degraders, i.e., scavengers. It is currently not clear how such remineralizing communities assemble at the microscale – how interactions between the different guilds influence their growth and spatial distribution, and hence the development and dynamics of the community. Here we address this knowledge gap by studying marine microbial communities that grow on the abundant marine biopolymer alginate. We used batch growth assays and microfluidics coupled to time-lapse microscopy to quantitatively investigate growth and spatial distribution of single cells. We found that the presence of exploiters or scavengers alters the spatial distribution of degrader cells. In general, exploiters and scavengers – which we collectively refer to as consumer cells – slowed down the growth of degrader cells. In addition, coexistence with consumers altered the production of the extracellular enzymes that breakdown polymers by degrader cells. Our findings reveal that ecological interactions by non-degrading community members have a profound impact on the functions of microbial communities that remineralize carbon biopolymers in nature.

**Importance:** Biopolymers are the most abundant source of carbon on the planet and their breakdown by microbial degraders releases metabolic products that allow cross-feeding cells to grow and fuel the assembly of microbial communities. While it is known that the growth of degraders can facilitate growth of downstream cross-feeders in microbial communities, it has remained generally unclear if and how cross-feeders influence growth of degraders. Bridging this knowledge gap is important because degraders primarily drive the remineralization of carbon, a central process in the carbon cycle. We found that the presence cross-feeders can influence the growth of degraders by altering their spatial distribution as well as extracellular breakdown enzyme activity. Our study sheds light on the role of microbial interactions in shaping the rate of carbon remineralization in nature.

## Introduction

Heterotrophic microbial communities drive the remineralization of carbon, which predominantly exist as biopolymers like chitin(1), alginate(2), cellulose(3) and xylan(4) in natural ecosystems, and thus drive a central step in the biogeochemical cycling of carbon(5–7). Within these communities, multiple microbial guilds coexist and engage in metabolic interactions. A key challenge of microbial ecology is to understand how these metabolic interactions influence the rate of carbon remineralization, which is an ecosystem function of major interest (8–11). The assembly of these communities follows intuitive rules (1). The first step in remineralization is carried out by specialized degraders that express and secrete hydrolytic enzymes that degrade biopolymers. The secretion of these enzymes leads to the formation of breakdown products. These products support the growth of the degraders and are also released into the environment, alongside with metabolites released by the growing degraders. The release of these metabolites creates niches for the growth of cross-feeder taxa that lack the ability to produce biopolymer-degrading enzymes but can utilize breakdown products (i.e. “exploiters”), or utilize other metabolites released from degraders and exploiter (i.e. “scavengers”; (9, 10, 12, 13)). Since degrader cells are positioned at the beginning of these food chains, their growth influences the assembly of downstream cross-feeders in biopolymer degrading communities(1, 10, 13). Therefore, any ecological interaction that impacts the growth of degrader cells is expected to impact the rate of carbon remineralization but the magnitude of these impacts is currently unknown.

Degrader cells often aggregate while growing on polysaccharides like chitin(14), alginate(15–17) or xylan(4, 18). Previous experimental and computational studies have established that aggregation, by increasing local cell density, allows degrader cells to benefit from the polysaccharide breakdown activities of neighboring cells(4, 15, 19). However, it is generally unclear how growth and collective behaviors of degraders are influenced by the coexistence with cross-feeding species. Since biopolymer breakdown and metabolic byproduct release at the microscale are a consequence of the activities of degraders, it is important to investigate the impacts of cross-feeding at the microscale on degrader taxa, in order to develop a better understanding on how interactions impact the growth of biopolymer degrading taxa and ultimately the rate of remineralization.

Here, we sought to address these knowledge gaps using simple two-species microbial communities that degrade alginate, an abundant biopolymer in marine ecosystems, on which the individual behavior of degraders is well characterized(20). We used a combination of batch growth assays and microfluidics coupled to time-lapse microscopy along with spatial analyses in order to quantitatively determine growth and aggregation behaviors of degrader cells when coexisting with cross-feeder cells. We find that the presence of cross-feeder cells influences the growth of degrader cells, indicating that the activity of cross-feeders can substantially alter the function of microbial communities.

## Results and Discussion

### Growth of communities composed of degrader and cross-feeder populations

As a first step, we measured the growth on alginate of three marine bacterial strains: i) *Vibrio cyclitrophicus* ZF270, ii) *Vibrio tasmaniensis* 1F187, and iii) *Ruegeria* sp. A3M17. Our goal was to investigate whether these natural marine isolates can be used to construct simple communities composed of two strains: an alginate degrader and a cross-feeder. *V. cyclitrophicus* ZF270 has the genetic repertoire to degrade polymeric alginate (alginate) resulting in production of oligo- or monomeric products of alginate breakdown (d-alginate)(2, 17, 21, 22). *V. tasmaniensis* 1F187 and *Ruegeria* sp. A3M17 cells lack alginate lyases but have a set of genes that could potentially confer the ability to import and utilize simpler oligo- or monomeric products resulting from alginate degradation(2, 10). In line with these predictions, we found that only ZF270 cells grew on alginate, whereas the cross-feeders 1F187 and A3M17 could not grow on alginate (Figure S1). We then grew strains on enzymatically hydrolyzed alginate, i.e. d-alginate, thus creating a scenario where cross-feeders experience byproducts released from alginate breakdown. We found that 1F187 grew to substantially higher levels on d-alginate compared to alginate, whereas A3M17 did not benefit from the hydrolysis products (Figure S1). These observations suggest that while 1F187 can indeed cross-feed on byproducts of alginate breakdown, A3M17 cannot cross-feed on breakdown byproducts despite encoding potential genes for uptake and utilization.

Based on these growth results we constructed simple two-species communities consisting of the degrader ZF270 and either the oligo-saccharide exploiter 1F187 or the byproduct scavenger A3M17 (Figure 1A). We then asked if presence of degraders allowed cross-feeders to grow. In addition, this growth assay enabled the quantification of any beneficial or detrimental effects of cross-feeder cells on degrader cells. When we grew degrader and cross-feeder cells in isolation or together on alginate in shaking flasks, we found that both cross-feeding strains 1F187 (Figure 1B) and A3M17 (Figure 1C) had increased growth yields in the presence of the degrader ZF270. The enhanced growth of A3M17 in coculture with degrader cells suggests that these cells likely cross-feed on metabolic byproducts that are secreted by ZF270 in coculture. Degrader cells displayed enhanced growth yields in the presence of 1F187 exploiter cells (Figure 1B) potentially through the removal of byproduct by the cross-feeding cells. In contrast, the growth yield of the degraders was similar in the presence or absence of A3M17 scavenger cells (Figure 1C), These observations indicate that, in addition to benefitting from the presence of degrader cells within communities, certain cross-feeder strains can also enhance the growth of degrader cells.

**Figure 1.**
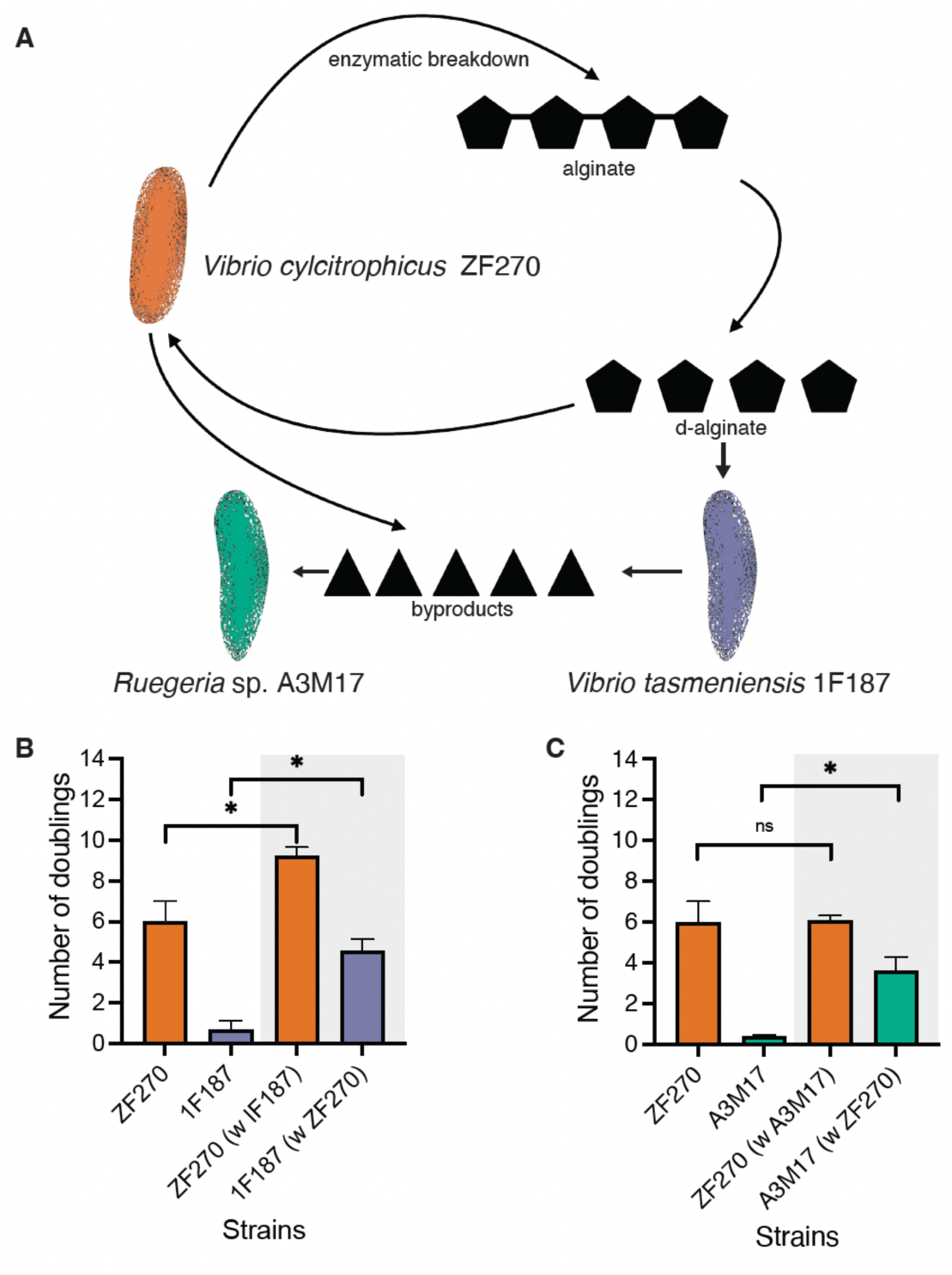
Growth dynamics of strains within simple two-species microbial communities in well-mixed environments. **(A)** *V. cyclitrophicus* ZF270 cells degrade the marine polysaccharide alginate, thereby producing oligo- and monomeric breakdown products(2, 17, 21, 22). Both *V. tasmaniensis* 1F187 and *Ruegeria* sp. A3M17 cells cannot grow on polymeric alginate (alginate), but 1F187 can act as an exploiter and grow on degradation products of alginate (d-alginate) while A3M17 can act as a scavenger by feeding on byproducts released from degrader cells (Figure S1). (**B** and **C**) The cross-feeders can grow in co-culture with the degraders. Growth yield represented as doublings. Growth was measured from colony counts obtained at the start and end of a growth cycle (see Methods). (**B**) *V. cyclitrophicus* ZF270 and *V. tasmaniensis* 1F187; or (**C**) *V. cyclitrophicus* ZF270 and *Ruegeria* sp. A3M17 in monocultures (no grey shading) or cocultures (grey shading) in shaking flasks with alginate. The bars indicate the mean whereas the error bars indicate the standard deviation (sd). Asterisks and ‘ns’ indicate statistically significant or non-significant comparisons, respectively, amongst groups (in B and C: paired t-tests: ZF270 vs ZF270 + 1F187: *p* =0.015, *t* =4.05, *R*^2^=0.80, *n_populations_*=3; ZF270 vs ZF270 + A3M17 : *p* =0.44, *t* =0.85, *R*^2^=0.15, *n_populations_* =3; 1F187 vs 1F187 + ZF270: *p* =0.013, *t* =4.25, *R*^2^=0.81, *n_populations_* =3; A3M17 vs A3M17 + ZF270 : *p* =0.04, *t* =5.95, *R*^2^=0.89, *n_populations_* =3).

### Cross-feeders influence growth of degraders through secreted compounds in model marine communities

A possible explanation for how cross-feeder cells influence the growth of degrader cells is through the production or consumption of secreted metabolites(11, 13, 23–27). Cross-feeders can decrease the growth of degraders by consuming metabolites that represent nutrients for degraders, thus reducing the availability of these for degraders, or by producing metabolites that inhibit the growth of degraders(26, 28). Conversely, cross-feeders can increase the growth of degraders by utilizing metabolites that inhibit the growth of degrader cells (13, 24) or by producing metabolites that support degrader growth. To understand the effect of compounds secreted by cross-feeders on the growth of degrader cells, we grew cross-feeder cells, harvested the spent media and grew degraders on this spent-media. Since cross-feeders do not grow on alginate, we grew them on substrates permissive to cross-feeder growth. Oligomer-exploiting 1F187 cells were grown on d-alginate for 36 h. A3M17 cells, which scavenge byproducts and did not grow on alginate or d-alginate (Figure S1), were grown on alginate supplemented with 0.1% marine broth for 36 h. The spent medium in either case was harvested to remove cross-feeder cells and used to monitor growth of degrader cells (Figure 2A and B). The growth of degraders on spent-medium was then compared to the growth of degraders on spent-medium produced by growing degrader cells on cognate fresh medium (Figure 2A and B). By comparing the growth of degraders on the spent medium of cross-feeders versus the spent medium of degraders, we were able to differentiate effects on the growth of degraders due to compounds released or utilized by cross-feeders from effects due to the consumption of limiting resources by the cross-feeders.

**Figure 2.**
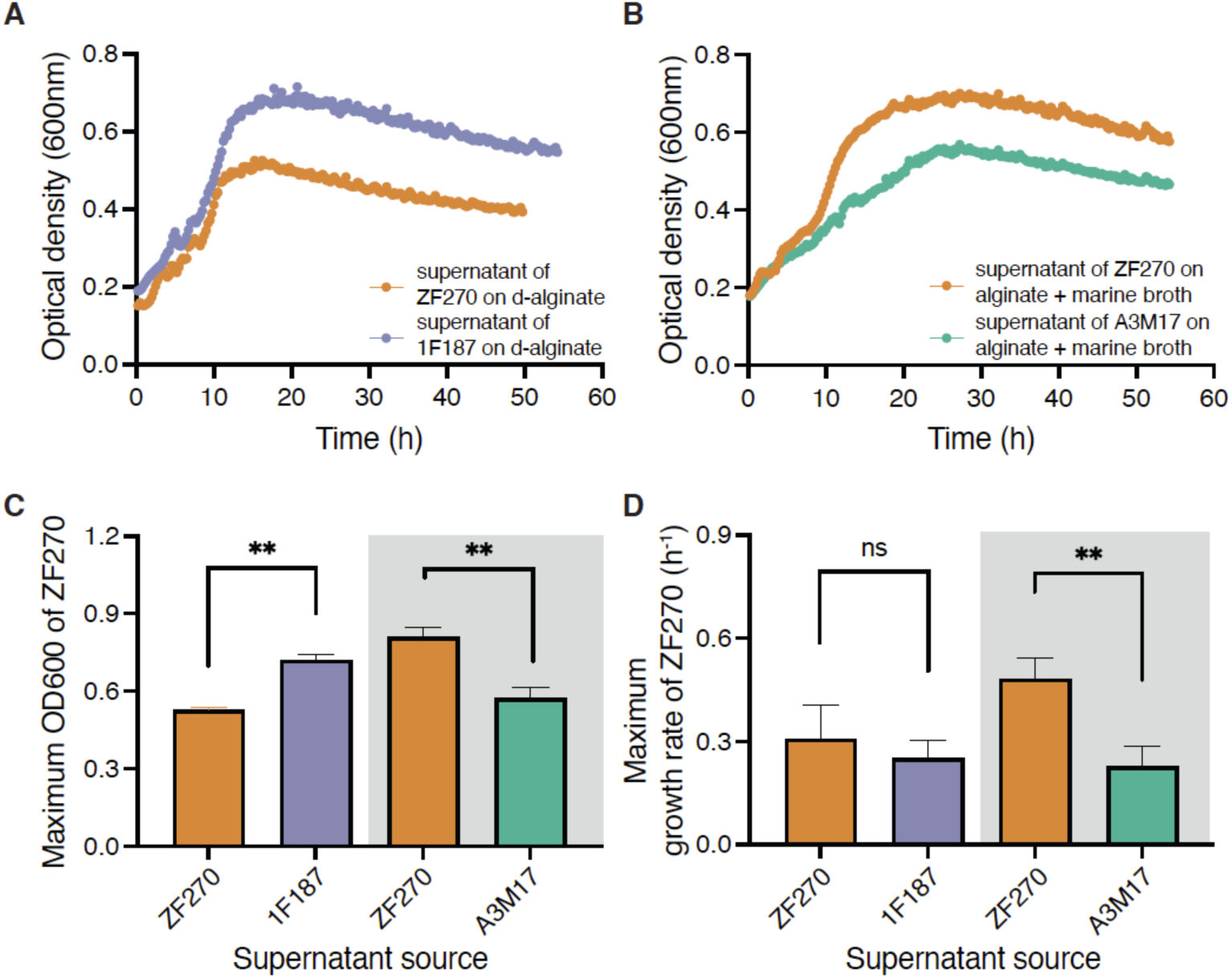
Spent-medium of cross-feeders alters growth dynamics of *V. cyclitrophicus* ZF270 degrader populations. (**A** and **B**) Growth dynamics (measured as optical density at 600 nm) of *V. cyclitrophicus* ZF270 cells in the presence or absence of spent-medium of (**A**) *V. tasmaniensis* 1F187 or (**B**) *Ruegeria* sp. A3M17. (**C**) Maximum population sizes and (D) maximum growth rates achieved by *V. cyclitrophicus* ZF270 cells in the various spent-media (ZF270: orange bars, 1F187: purple bars and A3M17: green bars). Asterisks and ‘ns’ indicate statistically significant and non-significant comparisons, respectively, amongst groups (in C: Mann-Whitney test, ZF270 and 1F187: *p* <0.0095, *n_populations_* = 4-6; ZF270 + A3M17: *p* <0.0043, *n_populations_* = 5-6; in D: Mann-Whitney test, ZF270 and 1F187: *p* <0.47, *n_populations_* = 4-6; ZF270 + A3M17: *p* <0.0043, *n_populations_* = 5-6).

We found that ZF270 degrader populations reached higher population sizes but showed similar growth rates on spent medium of 1F187 cells (Figure 2C and D) compared to spent medium of ZF270 cells. In contrast, ZF270 degrader populations had both reduced population sizes and growth rates on spent medium of A3M17 cells (Figure 2C and D) compared to spent medium of ZF270 cells. These observations indicate that cross-feeders indeed modify their external environment, through secretion or removal of metabolites and compounds that can positively (1F187) or negatively (A3M17) influence the growth of degrader cells. While the positive influence on ZF270 by 1F187 cells is in line with our earlier observation in cocultures (Figure 1B), the negative influence of A3M17 spent-medium on ZF270 cells (Figure 2C) is a deviation from the neutral effect observed in cocultures (Figure 1C).

### Presence of cross-feeders influences spatial distribution of degrader populations

Since interactions between cross-feeder and degrader cells often manifest at the microscale in natural ecosystems(9, 19, 29), we tested the influence of cross-feeder cells on degrader cells within microfluidic growth devices. Within these devices, cells can grow as monolayers inside growth chambers and move freely while receiving a constant supply of medium with alginate (4, 17, 18). Therefore, the number of cells and their positioning within microfluidic chambers is determined by the cellular growth rate as well as by cell movement(18). ZF270 cells are known to form aggregates when growing on alginate in microfluidic growth chambers(17) and the increased local cell density due to aggregation allows cells to cooperatively degrade alginate. In accordance with previous findings, we found that ZF270 cells form aggregates when growing on alginate (Figure 3A, Supplemental Video 1). In the presence of 1F187 cells (Figure 3B, Supplemental Video 2) the number of degrader cells within growth chambers was similar to that in monocultures of degraders (Figure 3D). In contrast, the number of degrader cells was reduced when A3M17 cells were present (Figure 3C and D, Supplemental Video 3). These results indicate that microscale interactions with cross-feeders can impact the growth rate or movement of degrader cells.

**Figure 3.**
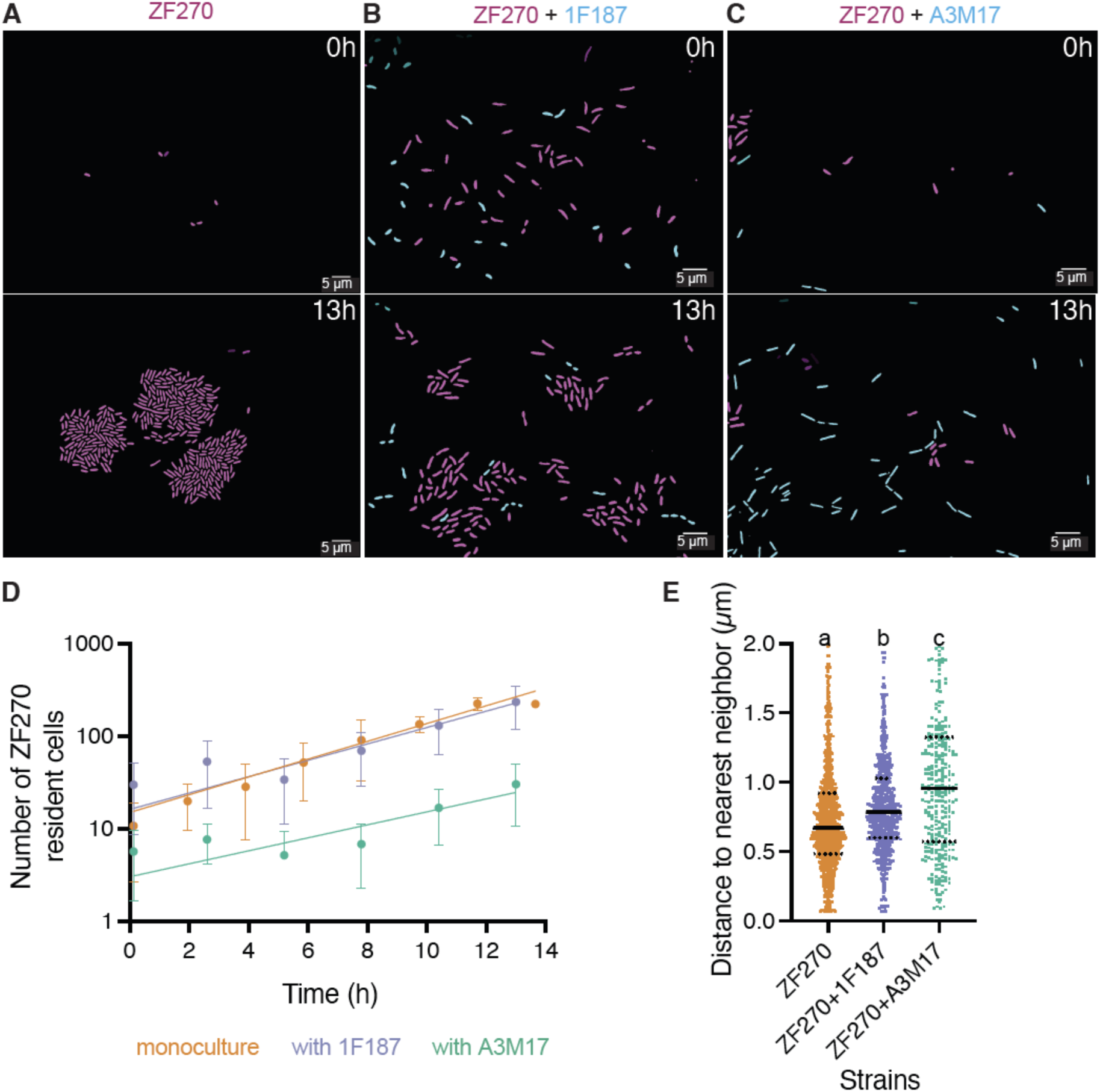
Aggregate formation by *V. cyclitrophicus* ZF270 populations is influenced by the presence of cross-feeders. Representative images of microfluidic growth chambers containing (**A**) monocultures of *V. cyclitrophicus* ZF270, (**B**) cocultures of *V. cyclitrophicus* ZF270 with *V. tasmaniensis* 1F187, and (**C**) *V. cyclitrophicus* ZF270 with *Ruegeria* sp. A3M17 cells. The upper panel are representative images of chambers at initiation of growth i.e. 0 h whereas the lower panels are images of chamber 13h after initiation. *V. cyclitrophicus* ZF270 are false colored in magenta whereas cells of *V. tasmaniensis* 1F187 and *Ruegeria* sp. A3M17 are false colored in cyan. (**D**) Some cross-feeders can influence group formation by degrader cells. Lines depict the number of cells present in the growth chamber over time. In the absence of cross-feeders (orange), or when present along with *V. tasmaniensis* 1F187 (purple), there is a higher number of *V. cyclitrophicus* ZF270 resident cells over time within chambers compared to cocultures with *Ruegeria* sp. A3M17 (green). Circles indicate the number of cells present at a given time point in each replicate chamber (*n*_chambers_ = 8-9). Lines are fits of a mean linear regression line for each condition – ZF270: *R*^2^ = 0.75 and *k* = 16.86h^-1^; ZF270+1F187: *R*^2^ = 0.42 and *k* = 13.95 h^-1^; ZF270+A3M17: *R*^2^ = 0.32 and *k* = 1.41h^-1^. Line depicting ZF270+A3M17 is statistically different from the line showing the trajectory of ZF270 monocultures (Statistically different slopes of the linear regressions: *F* = 97.11. DF_n_ = 1, DF_d_ = 81, *p*<0.0001). Lines depicting trajectories of ZF270 monocultures and ZF270+1F187 are statistically similar (*F* = 1.047. DF_n_ = 1, DF_d_ = 94, *p*=0.30). (**E**) Nearest neighbor distances of every ZF270 cell in monocultures (orange) or coculture with 1F187 (purple) or A3M17 (green). Violin plots indicate the nearest neighbor distances of ZF270 cells in microfluidic growth chambers (*n*_chambers_ = 8-9). Asterisks indicate statistical differences between groups based on a One-way ANOVA and Dunn’s post-hoc test (*p* <0.045 and statistic 10.81). Bold black lines indicate the median while dashed black lines indicate the 25^th^ and 75^th^ quantiles of the distribution. Violin plots are scaled by width.

To understand how oligomer-exploiters and by-product scavengers influence the spatial distribution of degrader cells, we quantified the intercellular distances between degrader cells in the absence or presence of one the cross-feeder strains. For this, we utilized the nearest neighbor approach and quantified the intercellular distance between a focal ZF270 cell and its nearest neighbor in each growth chamber (Figure 3E). This analysis revealed that compared to monocultures, the presence of 1F187 cells resulted in a statistically significant increased intercellular distances of degrader cells by about 8% (Figure 3B and E), whereas the presence of A3M17 cells resulted in a statistically significant increased intercellular distances of degrader cells to a higher degree, by about 58% (Figure 3C and E). These results indicate that the presence of either type of cross-feeder strain can alter the spatial distribution of degrader cells. In addition, the detrimental effect of A3M17 scavengers observed in well-mixed experiments (Fig. 2C) is in line with the reduction in group sizes (Fig. 3D) and altered spatial distribution (Fig. 3E) of degraders at the microscale.

### Coexistence with cross-feeding cells shifts the temporal growth dynamics of degrader cells

Since the presence of cross-feeders alters the spatial behaviors of degrader cells, and the growth of degrader cells on polymers depends on increased cell densities(4, 17), we expected interactions with cross-feeders to alter growth dynamics of degraders. To quantify these effects, we measured the single-cell growth rates of ZF270 cells that were grown on alginate in the absence or presence of either cross-feeding cell-type (Figure 3). For this, we used image analysis to segment and track cells and we mapped the lineages of all cells present in each growth chamber based on division events. This analysis allowed to quantify the rate at which individual cells in a microfluidics growth chamber increase their length and divide. We then analyzed the distribution of growth rates of single cells that were born at different time points in microfluidic growth chambers. We found that degrader ZF270 cells growing in monoculture achieved their maximum growth rates 8 h after the start of the growth experiment (Figure 4). However, the presence of either cross-feeder reduced the maximum median growth rate of degrader cells and increased the time to reach the maximum growth rate. When oligomer-exploiter 1F187 cells were present, ZF270 cells reached their maximum growth rate after approximately 10 h of growth initiation, whereas scavenger A3M17 cells delayed the time to maximum growth rate of degrader cells to 13 h (Figure 4).

**Figure 4.**
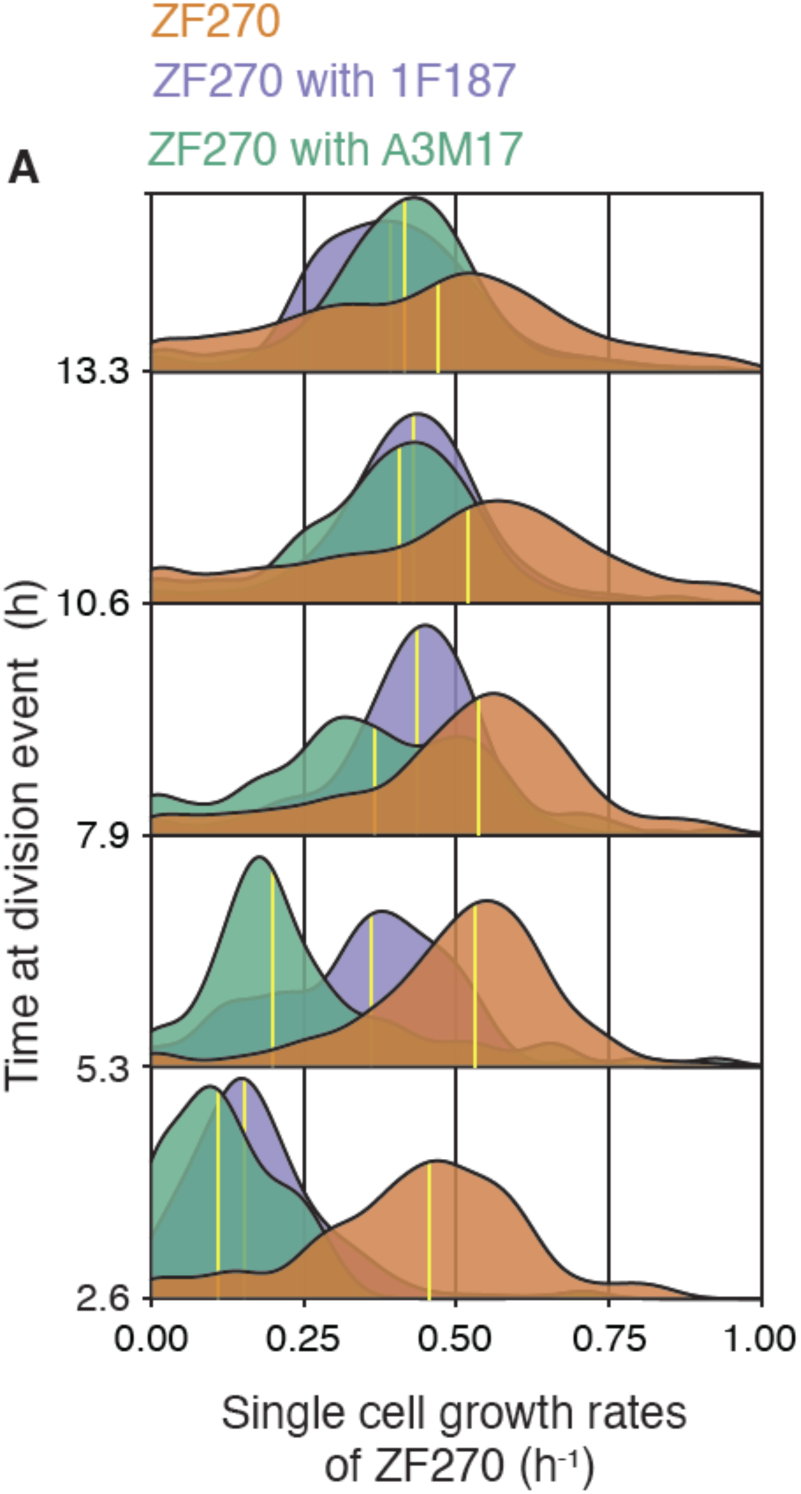
Presence of cross-feeder cells shifts the temporal growth dynamics of degrader cells. Single-cell growth rates of all *V. cyclitrophicus* ZF270 cells growing in microfluidic growth chambers were measured in monocultures (orange) or in cocultures with *V. tasmaniensis* 1F187 (purple) and *Ruegeria* sp. A3M17 (green) cells. Growth rates of cells were binned into time intervals based on the time they emerged from division. ZF270 cells growing in monoculture reach maximal growth rates faster than *V. cyclitrophicus* ZF270 cells growing along with *V. tasmaniensis* 1F187 and *Ruegeria* sp. A3M17.The distributions are pooled for all replicates. Cells were binned into 2.65h intervals based on the time at a division event (bins: 0-2.65h, 2.66-5.30h, 5.31-7.9h, 7.91-10.60h and 10.61-13.4h). Yellow horizontal lines depict the medians of the distributions (n_cells_ of ZF270: monocultures=10585, with 1F187=7698, with A3M17=1485). Medians of individual chambers are statistically distinct. (One way ANOVA; 2.6h: *R*^2^= 0.95, *p* <0.0001, *F* = 212.7; 7.9h: *R*^2^= 0.42, *p* = 0.0025, *F* = 7.99)

**Figure 5.**
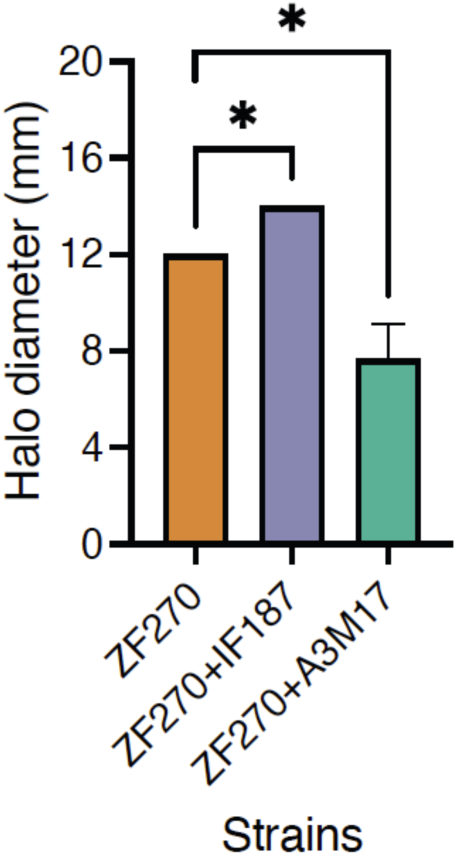
Cross-feeders alter alginate lyase secretion by degraders. *V. cyclitrophicus* ZF270 cells have increased secretions of alginate lyases when growing together with *V. tasmaniensis* 1F187 cells and decreased secretion of alginate lyases when growing along with *Ruegeria* sp. A3M17 cells compared to ZF270 cells in monoculture. Alginate lyase secretion is measured using an iodine assay after an 18-hour growth cycle. The diameter of the halo (mm) is used as a proxy for alginate lyase production (see Materials and Methods). The bars represent the mean of the halo diameter (*n* = 3) while error bars indicate the 95% confidence intervals (CI). Asterisks indicate statistically significant differences between comparisons (One-way ANOVA, *F* = 283, *p* = 0.0035, *R*^2^= 0.99).

The detrimental effect of 1F187 on the growth dynamics of the degraders (Figure 4) was in contrast to the observation in well-mixed environments that spent-medium of 1F187 cells benefited ZF270 cells (Figure 2A and C). Such an observation could arise if 1F187 cells compete with the degrader cells at the microscale for breakdown products resulting from alginate degradation. The findings that byproduct scavenging A3M17 cells increased both intercellular distances (Figure 3E) and growth rates (Figure 4) of ZF270 cells, align well with the observations of the spent-medium experiments, further lending credence to the idea that A3M17 cells negatively influence the growth of degrader cells, potentially through secretion or utilization of compounds. In addition, since growth of ZF270 cells was dependent on cell density(17), a reduced density in the presence of A3M17 cells likely causes ZF270 cells to have reduced growth rates. When the growth of cross-feeders was analyzed, the byproduct scavenging A3M17 achieved higher growth rates compared to the oligomer-exploiter 1F187 (at 7.9 h, Figure S3). Taken together, our results suggest that the presence of cross-feeders alters growth dynamics of degrader cells at the microscale.

### Cross-feeder cells alter activity and secretion of polysaccharide degrading enzymes by degraders

Since the presence of cross-feeders alters the growth dynamics of degrader cells it is plausible that the activity of alginate lyases, enzymes that mediate breakdown of alginate, is also altered. In ZF270 cells, alginate lyases are secreted extracellularly(17). Therefore, quantifying the activity of alginate lyases enabled the measurement of the effect of cross-feeders on the enzyme activity of degraders. For this, we used a plate-based assay where ZF270 monocultures or cocultures were spotted on alginate plates along with one of the cross-feeders. In this assay, alginate reacts with iodine to produce a violet compound. The breakdown of alginate by secreted enzymes that diffuse away from the cells creates simpler oligomers or monomers that yield colorless halos when stained with iodine. We measured the diameter of the halo as a proxy for understanding activity of ZF270 cells in the absence or presence of degraders.

We found that ZF270 cells in coculture with 1F187 cells had higher alginate lyase activity than ZF270 cells in monoculture. A similar observation has been made in the case of increased expression of chitinases by *Vibrio natrigens* in response to cross-feeding by *Alteromonas macleodii* cells(27). We speculate that increased alginate lyase activity combined with competition for oligomeric breakdown products between ZF270 and 1F187 cells causes a shift in the growth dynamics of ZF270 cells (Figure 4).

In contrast, ZF270 cells in coculture with A3M17 cells had reduced lyase activity compared to ZF270 cells in monocultures. Since A3M17 cells negatively influence the growth of ZF270 cells (Figure 2B), reduced alginate lyase activity can likely be attributed to reduced growth of ZF270 cells. Taken together, our findings indicate that in addition to benefitting from the metabolic activities of degrader cells, the by-product cross feeding of A3M17 cells influences the growth dynamics of degrader cells. The presence of A3M17 alters the activity of polysaccharide degrading enzymes as well as the spatial organization of degrader cells.

Our findings indicate that cross-feeding, which is a prevalent phenomenon in natural microbial communities(11), can alter the growth of species that remineralize carbon biopolymers. It is known that biopolymer degrading bacteria can alter their surface association in response to changes in environmental nutrient composition(4, 22, 30). Our findings indicate that differences in collective behaviors including the spatial distribution of cells can also arise as a consequence of species interactions. These observations suggest that degraders not only structure microbial food-chains but their activity is also shaped by downstream cross-feeding species.

Growth dynamics and behaviors such as aggregation or dispersal are usually studied in clonal populations. Similarly, degradation abilities of species are studied in monoculture experiments. Our findings reveal the importance of understanding such behaviors in the context of a multi-species community because members downstream of the actual degradation influence growth of degrading strains. In addition, our study demonstrates that the type of cross-feeding interaction determines the impact on growth of taxa that form the base of polysaccharide degrading food webs. This finding highlights the necessity of taking into account ecological interactions in addition to *in silico* predictions of ecosystem function based on the enzymatic functions encoded within metagenomes(31). Ultimately, our results underscore the importance of understanding behaviors at the level of single cells(32) and investigating how individuals within communities react to the presence of cells belonging to different species. Uncovering the molecular mechanisms that underlie behavioral responses will enable a stronger understanding of not only the drivers of community assembly and spatial organization within communities but also allow the development of potential applications to systematically control the composition and behavior of microbial ecosystems.

## Materials and Methods

### Bacterial strains, media and growth assays

We used *Vibrio cyclitrophicus* ZF270, *Vibrio tasmaniensis* 1F187 and *Ruegeria* sp. A3M17 that contained plasmids bearing fluorescent proteins: ZF270 – pLL103-mKate2, 1F187 – pLL102-mCitrine and A3M17-pJES005-mTurq. Plasmids pLL103-mKate2 and pLL102-mCitrine were constructed as described previously(33). Plasmid pJES005-mTurq was created by modifying the pBBR1MCS mCherry plasmid pGS001(34). Plasmids pGS001 and pOXC101(33) were extracted from *Escherichia coli* DH5α and β3914 respectively using the QIAprep Spin Miniprep Kit (Qiagen). A tac promoter, lac operator, and mCitrine fluorescent protein were amplified from plasmid pOXC101 (Pollak et al 2021) with the Phusion High-Fidelity PCR Master Mix (ThermoFisher) using primers JES003 (ATATCTCGAGGGCGCGCCTTGACAATTAATC) and JES004 (ATATTCTAGAGGATCCTTACTTATATAATTC). The PCR product was purified (QIAquick PCR Purification Kit; Qiagen) and digested alongside pGS001 for 2 hours at 37°C using XhoI and XbaI (New England Biolabs). Digested plasmid backbone and PCR product were gel extracted (QIAquick Gel Extraction Kit; Qiagen) and ligated for 2h at 24°C using T4 DNA ligase (Invitrogen). The newly ligated plasmid pJES002 was transformed into chemically competent *E. coli* Stellar cells (Clontech) and plated onto Luria Bertani agar supplemented with 25 µg/ml chloramphenicol. Extracted pJES002 and pOXC102 (33) were digested using AscI and BamHI-HF (New England Biolabs), and the plasmid backbone of pJES002 and a tac promoter, lac operator, and mTurquoise2 fluorescent protein were gel extracted, ligated, and transformed as above. The generated plasmid pJES005 was confirmed via Sanger sequencing and finally electroporated into the conjugative strain *E. coli* RHO3+.

Strains were cultured in Marine Broth (DIFCO) and grown for 18 hours at 25 °C. Cells from these cultures were used for growth experiments in Tibbles Rawling (TR) minimal medium <u>[1</u>] containing either 0.1% (weight/volume) Polymeric alginate (Sigma Aldrich) or 0.1% degraded (d) alginate. D-alginate was produced by enzymatically degrading 2% alginate with 1 unit ml^-1^ of alginate lyase (Sigma Aldrich) at 37 °C for 24 hours. Carbon sources were prepared in nanopure water and filter sterilized using 0.40 μm Surfactant-Free Cellulose Acetate filters (Corning, USA). Well-mixed batch experiments in alginate or d-alginate media were performed in 96-well plates and growth dynamics were measured using a micro-well plate reader (Biotek, USA).

### Coculture growth assays

Coculture assays were initiated as described previously (36). To measure growth of degrader and cross-feeding strains in communities, ∼10^5^ colony forming units (CFUs) ml^−1^ of either strain were inoculated into 10 mL TR medium with the requisite carbon source and cell numbers were determined at 0h and 36h by plating. Strains display distinct morphologies and harbor different fluorescent phenotypic markers and therefore can be differentiated on Marine Agar (MB broth with 1.5% Agar (Applichem) plates. Growth of strains was determined by calculating the number of doublings of individual strains in mono- or co-cultures: Doublings = (log_2_ (*N_f_*/*N_i_*)), where *N_i_* is initial number of CFUs at 0h and *N_f_* is the final CFU count (36, 37).

### Spent-medium experiments

To generate spent-medium of degraders, *Vibrio cyclitrophicus* ZF270 cells (∼10^5^ CFUs ml^−1^) were grown in 10 mL TR medium with 0.1% alginate for 36 hours. Cells were separated from the supernatant by centrifugation at 10,000 rpm for 10 minutes followed by filtering through 0.40 μm Surfactant-Free Cellulose Acetate filters (Corning, USA). The supernatants were then used to grow cross-feeder populations, whose growth was initiated at starting densities of ∼10^5^ CFUs ml^−1^. To generate supernatants of *Vibrio tasmaniensis* 1F187, cells (∼10^5^ CFUs ml^−1^) were first grown on 0.1% d-alginate TR medium for 36 hours following which supernatants were harvested as described above. Growth of *Vibrio cyclitrophicus* ZF270 cells was then initiated at ∼10^5^ CFUs ml^−1^ on supernatant as well as on supernatant of d-alginate TR medium where ZF270 cells were grown as a control. To generate supernatants of *Ruegeria* sp. A3M17, cells (∼10^5^ CFUs ml^−1^) were grown on MB medium with 0.1% p-alginate and supernatants harvested as described above. Growth of *Vibrio cyclitrophicus* ZF270 cells was then initiated at ∼10^5^ CFUs ml^−1^ on supernatant as well as on supernatant of MB medium with 0.1% alginate where ZF270 cells were grown as a control.

### Microfluidics and time-lapse microscopy

Microfluidics experiments and microscopy were performed as described previously (4, 5, 38). Cell growth and behavior was imaged within chambers which ranged from 60-120 × 60 x 0.56 μm. (*l x b x h*) Within these chambers, cells attach to the glass surface and experience the medium that the diffused through the lateral flow channels. Imaging was performed using IX83 inverted microscope systems (Olympus, Japan) with automated stage controller (Marzhauser Wetzlar, Germany), shutter, and laser-based autofocus system (Olympus ZDC 2). Chambers were imaged in parallel on the same PDMS chip, and phase-contrast and fluorescent (mKate2 and/or mCitrine) images of each position were taken every 8 or 10 min. The microscopy unit and PDMS chip was maintained at 25 °C using a cellVivo microscope incubation system (Pecon GmbH, Germany).

### Alginate Lyase assay

We adapted a previously described agarose plate based assay(39) to test the ability of monocultures and cocultures to secrete alginate lyases. For each strain, cultures were growth for 18h in MB medium and 1 ml of cell suspension was centrifuged (13000 rpm for 2mins) in a 2ml microfuge tube. The supernatant was discarded and the cell-pellet was subject to two rounds of washing with TR medium without any carbon source. The cell pellet was suspended in 1ml of TR medium without carbon source and the optical density measured and adjusted to 0.1OD. For ZF270 monocultures 50µl of 0.1OD culture, or for cocultures 50μl of degrader and 50µl of cross-feeder cultures were mixed and spotted on plates that were made using TR medium containing 0.1% (w/v) alginate (Sigma Aldrich) and 1% agarose (Applichem). Colonies were allowed to grow for 30h at 25°C and then the plates were flooded with 2%. Gram’s Iodine (Sigma Aldrich). The excess iodine was discarded and the imaged using an iPhone 12 camera. If cells secreted alginate lyases, then a distinct clearance zone was formed, the diameter of which was measured using a standard ruler.

### Image analysis

Cells within microscopy mages were segmented using a custom built Python based (v3.7) segmentation workflow and tracked with SuperSegger(40). The output of segmentation and tracking were processed using Matlab v2017b or newer and R v4. Phase contrast channel images were used for alignment following which fluorescence channel images were used for segmentation, tracking and linking. Images were cropped at the boundaries of each microfluidic chamber. Growth properties and spatial locations were directly derived from the downstream processing tools of SuperSegger (gateTool and superSeggerViewer). Spatial distances between cells (Fig. 3E) were computed from segmentation data using the R package *spatstat*(41).

### Datasets and statistical analysis

All batch experiments were replicated 3-6 times. Growth curves were analyzed in Python v3.7 using the *Amiga* package(42) and GraphPad Prism v8 (GraphPad Software, USA). The microscopy dataset set consists of eight, nine and eight chambers, respectively for ZF270, ZF270+1F187 and ZF270 + A3M17. These are grouped into three biological replicates wherein each biological replicate is fed by media through a unique channel in a microfluidic chip. Cells with negative growth rates were excluded from the analysis after visual curation, and represent artefacts, mistakes in linking during the segmentation or tracking process or non-growing deformed cells. Each chamber was treated as an independent. Each figure depicts means or medians of all chambers for each condition. Pre-existing linear or exponential regression models in GraphPad Prism v 8.0 (GraphPad Software, USA) were applied to determine relationships between independent measures such as: number of cells versus time (Fig. 3D) and intercellular distance versus number of nearest neighbors (Fig. 3E). Comparisons were considered statistically significant when *P < 0.05* or when False Discovery Rate (FDR) corrected *Q < 0.05*. FDR corrections were applied when multiple t tests were performed for the same dataset. Measures of effect size are represented by the R^2^ or eta^2^ value. All statistical analyses were performed in GraphPad Prism v 8.0 (GraphPad Software, USA) and Rstudio v1.1.463 (Rstudio inc).

## Author Contributions

GD, JS, and JK conceived the research along with OC, RS and MA. JS and JES constructed the plasmids and strains. GD performed all experiments. MD performed pilot experiments. GD analyzed the data with inputs from JS, JK and MA. GD wrote the manuscript with inputs from JS, JK, RS and MA.

## Acknowledgements

We thank past and present members of the Microbial Systems Ecology group for feedback. This research was funded by an ETH fellowship and a Marie Curie Actions for People COFUND program fellowship (FEL-37-16-1) to GD; the Simons Foundation Collaboration on Principles of Microbial Ecosystems to OC, RS and MA (PriME #542379, #542383 and #542395); and by ETH Zurich and Eawag.

## Data Availability

All raw curated image analysis datasets and source data for figures will be deposited in the ERIC and Zenodo repositories upon publication. Supplemental Videos are available on FigShare (doi: https://doi.org/10.6084/m9.figshare.22317004.v1)

