## Supplemental information for "Interspecies interactions determine growth dynamics of biopolymer degrading populations in microbial communities"

[1]Microbial Systems Ecology Group, Department of Environmental Systems Sciences, Institute of Biogeochemistry and Pollutant Dynamics, ETH-Zurich, 8006 Zurich, Switzerland
[2]Department of Environmental Microbiology, Eawag: Swiss Federal Institute of Aquatic Sciences, 8600 Duebendorf, Switzerland
[3]Department of Civil and Environmental Engineering, MIT, Cambridge, MA 02139, USA
[4]Current address: Department of Biology, University of Southern California, Los Angeles, CA 90089
[5]Institute of Environmental Engineering, Department of Civil, Environmental and Geomatic Engineering, ETH Zurich, 8093 Zurich, Switzerland
[6]Department of Marine Sciences, University of Georgia, Athens, GA 30602
[7]Environmental Engineering Institute, EPFL, Lausanne, Switzerland

Classification: Biological Sciences/Microbiology

Keywords: cross-feeding, aggregation, community assembly, dispersal, spatial organization, biopolymer degradation

**This file includes:**

Supplementary Figures 1 to 3
Supplementary Videos 1 to 3
Supplementary Text

**Supplemental Figures:**

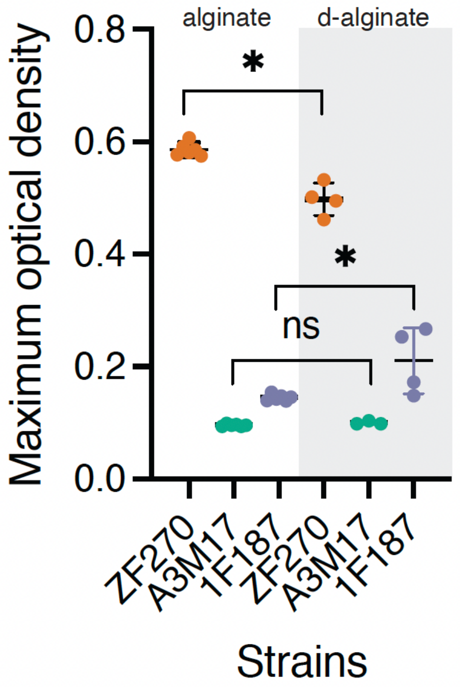


**Figure S1. Growth of strains on alginate and d-alginate.** ZF270 cells can degrade alginate and hence can grow on polymeric alginate (alginate, white panel) whereas cross-feeders cannot grow substantially on alginate. Growth of 1F187 cells is substantially increased on d-alginate whereas A3M17 do not grow on either carbon source. Shown is maximum optical density (OD) achieved by *V. cyclitrophicus* ZF270 (orange dots), *V. tasmaniensis* 1F187 (purple dots) and *Ruegeria* sp. A3M17 (green dots) populations over the course of a 40h growth cycle. Horizontal lines represent the mean maximum OD ($n$ = 3-6) while error bars indicate standard deviation (sd).

Asterisks or ns indicate statistically significant or non-significant comparisons, respectively amongst groups (Mann-Whitney tests: ZF270 – *p*=0.0095, n=6-4; A3M17- *p*=0.08, n=3-6; 1F187 - *p*=0.01, n=3-6).

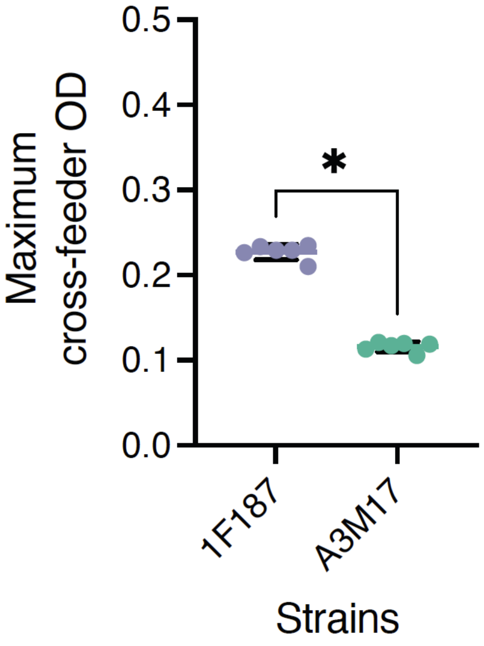


**Figure S2. Spent medium of *V. cyclitrophicus* ZF270 populations can support the growth of oligo-saccharide cross-feeders.** *V. tasmaniensis* 1F187 (purple dots) populations reach higher maximum ODs compared to *Ruegeria* sp. A3M17 (green dots) populations on spent-medium of *V. cyclitrophicus* ZF270 populations. Maximum optical density (OD) achieved by (orange dots), and over the course of a 40h growth cycle. Horizontal lines represent the mean maximum OD ($n$ = 3) while error bars indicate 95% confidence intervals (CI). Asterisks or ns indicate statistically significant or non-significant comparisons, respectively amongst groups (Independent sample t-tests: $p < 0.0001$, t = 25.89, $R^2$ = 0.98, n = 6)

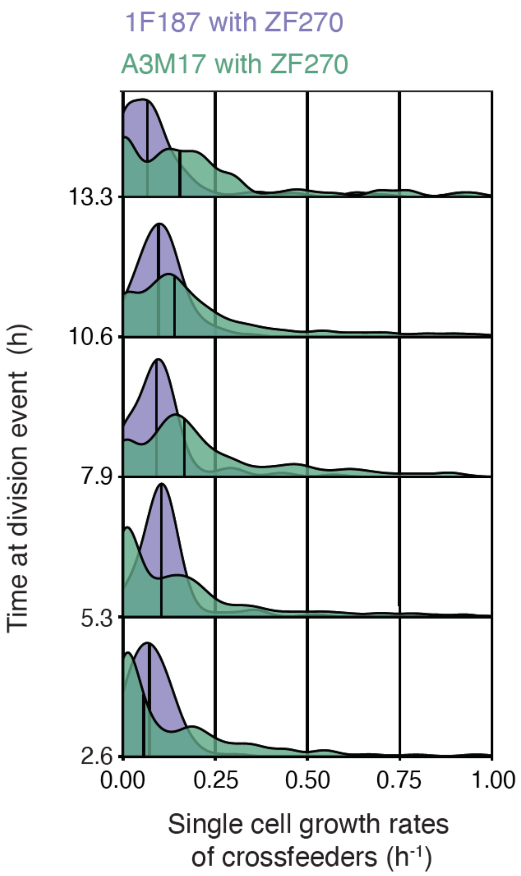


**Figure S3. Single cell growth rate distributions of crossfeeder cells when growing along with *V. cyclitrophicus* ZF270 cells.** Time-binned single cell growth rate distributions of *V. tasmaniensis* 1F187 (purple) and *Ruegeria* sp. A3M17 (green). The distributions are pooled for all replicates. Cells were binned into 2.65-hour intervals based on the time at division (bins: 0-2.65h, 2.66-5.30h, 5.31-7.9h, 7.91-10.60h and 10.61-13.4h). Yellow horizontal lines depict the medians of the distributions ($n_{cells}$ of cross-feeders: 1F187 = 1499, with A3M17 = 5521). Medians are statistically distinct at 7.9h. (Independent samples t-test; 7.9h: $R^2 = 0.95$, $p = 0.0005$, $t = 4.63$; $df = 13$, 7.9h: $R^2 = 0.62$)

**Supplementary videos:**

Cells growing in microfluidic growth chambers. Cells are false colored based on expression of GFP by ZF270 cells or mCitrine by 1F187 or A3M17 cells. Frame rate for all movies is 20 images per second. Scale bar is 5 microns. Available on FigShare; DOI: https://doi.org/10.6084/m9.figshare.22317004.v1

**Supplemental Video 1**. Time-lapse of *Vibrio splendidus* ZF270 (false coloured in magenta) cells within a representative microfluidics chamber fed with 0.1% alginate as the sole carbon source.

**Supplemental Video 2**. Time-lapse of *Vibrio splendidus* ZF270 cells (false coloured in magenta) growing in the presence of *V. tasmaniensis* 1F187 (false coloured in cyan) within a representative microfluidics chamber fed with 0.1% alginate as the sole carbon source.

**Supplemental Video 3**. Time-lapse of *Vibrio splendidus* ZF270 cells (false coloured in magenta) growing in the presence of *Ruegeria* sp. A3M17 (false coloured in cyan) within a representative microfluidics chamber fed with 0.1% alginate as the sole carbon source.
